# Pan-B-Lineage Targeting with Adapter CAR-T Cells Controls Antigen-Heterogeneous Lymphoma

**DOI:** 10.64898/2026.09.12.751171

**Authors:** Simon Krost, Anna-Sophia Mast, Daniel Atar, Beate Kristmann, Sophia Scheuermann, Christian M. Seitz

**Author notes:** Corresponding Author: **Christian M. Seitz,** Department of General Pediatrics, Hematology and Oncology, University Children’s Hospital, Tuebingen, Germany.

## Abstract

Antigen heterogeneity and antigen-negative relapse represent major limitations to durable CAR-T cell efficacy in B-lineage malignancies. We previously developed the Adapter CAR-T cell (AdCAR-T) platform, which enables flexible redirection of engineered T cells to distinct surface antigens through biotinylated adapter molecules (AMs). In this study, AMs generated from an in-house-produced tafasitamab biosimilar (anti-CD19), commercial rituximab (anti-CD20), and an in-house-produced daratumumab biosimilar (anti-CD38) mediated potent, antigen-specific AdCAR-T cell cytotoxicity. While single-antigen targeting resulted in the selection of antigen-negative tumor populations, simultaneous targeting of CD19, CD20, and CD38 effectively controlled a defined heterogeneous Burkitt lymphoma model *in vitro* and induced sustained tumor control *in vivo*. Selective loss of the CD38^+^ AdCAR-T cell population after CD38-directed AM exposure was consistent with fratricide; however, the surviving CD38^low^ population retained cytotoxic activity. These findings establish combinatorial AdCAR-T cell targeting as a flexible pan-B-lineage strategy for addressing pre-existing antigen heterogeneity and support further development of antibody-derived AM combinations for B-cell malignancies.

## Introduction

Chimeric antigen receptor (CAR)-T cell therapy has substantially improved outcomes for patients with relapsed or refractory B-lineage malignancies in the last decades (1–3). However, relapse due to loss or downregulation of the targeted antigen remains a major challenge in antigen-directed immunotherapy (4–8). Therapeutic immune pressure can select pre-existing antigen-deficient tumor cell clones or promote genetic and phenotypic alterations that reduce antigen expression and impair target recognition. Addressing both interpatient and intratumoral antigen heterogeneity is therefore a central objective in the development of more durable CAR-T cell therapies.

Several approaches have been developed to enable multi-antigen tumor recognition, including dual- and trispecific CAR constructs and the coadministration of separately manufactured CAR-T cell products (9–11). Although these strategies broaden antigen recognition, their target sets are predefined before infusion. The Adapter CAR-T cell (AdCAR-T) platform separates antigen recognition from cellular signaling: AdCAR-T cells recognize a universal linker-label epitope (LLE) on biotinylated adapter molecules (AMs), which in turn bind their respective tumor antigens (Figure 1A). Antigen recognition can thereby be reconfigured after AdCAR-T cell infusion, and therapeutic activity adjusted through AM selection, dosing, and scheduling (12–15). Previous AdCAR-T studies from our group established flexible targeting across models of B-lineage lymphoma, acute myeloid leukemia (AML), breast cancer, and small cell lung cancer, and demonstrated that combinatorial targeting can prevent antigen escape in AML (12–15). Consistent findings were subsequently reported using another adapter-CAR platform in heterogeneous AML (16). However, the efficacy of triple-AM targeting in a defined antigen-heterogeneous B-cell lymphoma model *in vivo*, as well as the effect of CD38-directed targeting on the AdCAR-T cell compartment, remain unexplored.

**Figure 1.**
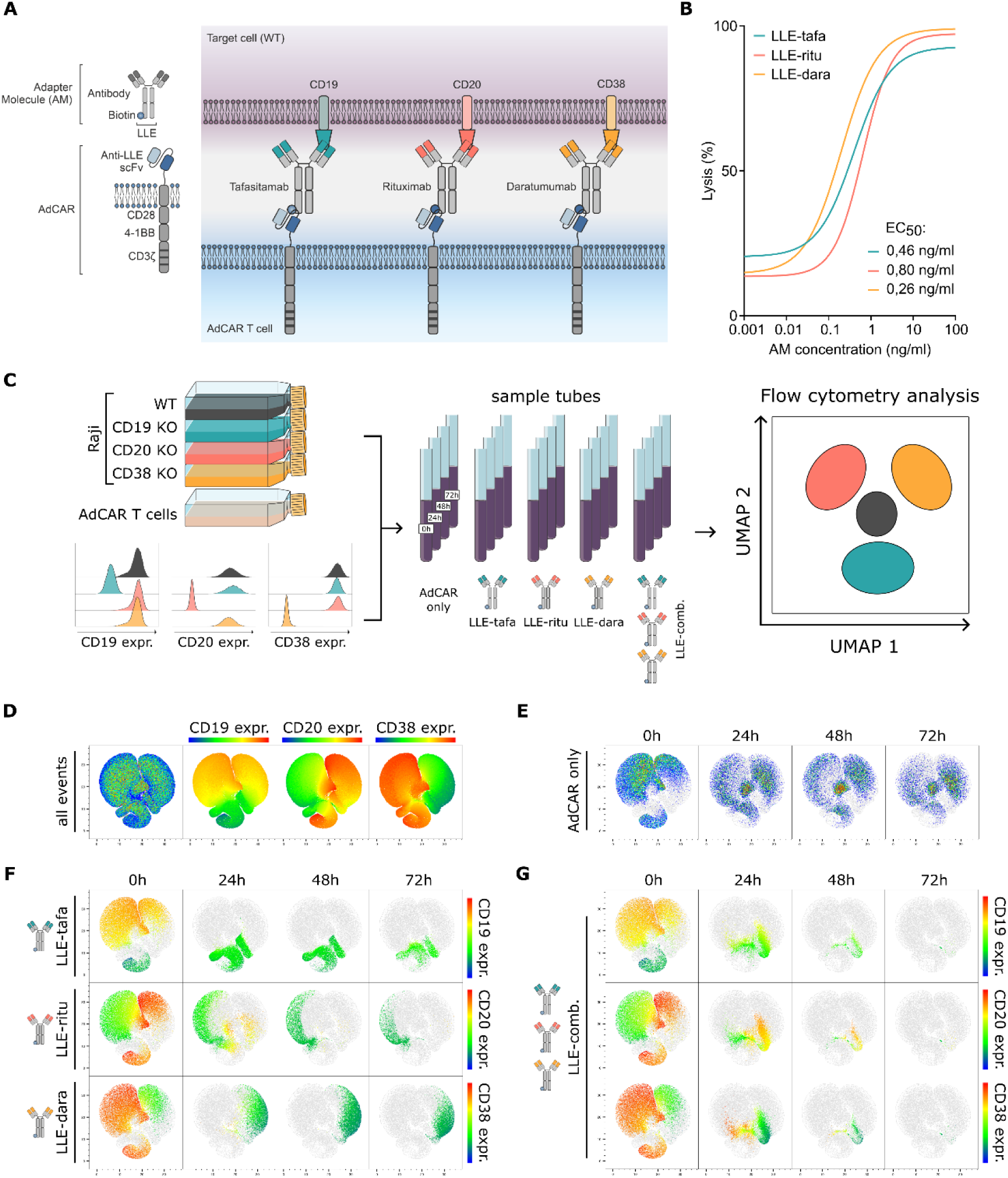
CD19-, CD20-, and CD38-directed AMs mediate antigen-specific, dose-dependent, and combinatorial AdCAR-T cell activity *in vitro*. **A**, Schematic of the AdCAR-T cell system. An in-house-produced tafasitamab biosimilar (anti-CD19), commercial rituximab (anti-CD20), and an in-house-produced daratumumab biosimilar (anti-CD38) were biotinylated to generate LLE-containing AMs that redirect AdCAR-T cells to the corresponding antigen. **B**, Lysis of Raji^WT^ cells after 72 hours at an E:T ratio of 4:1 across the indicated AM concentrations using AdCAR-T cells from one donor (six technical replicates per concentration). Curves show nonlinear fits of log-transformed concentration-response data. EC₅₀ values are shown; R² values were 0.9956 for LLE-tafasitamab, 0.9969 for LLE-rituximab, and 0.9918 for LLE-daratumumab. **C**, Experimental workflow. Equal numbers of Raji^WT^, Raji^CD19KO^, Raji^CD20KO^, and Raji^CD38KO^ cells were co-cultured with AdCAR-T cells under the indicated AM conditions for 0–72 hours before flow-cytometric and UMAP analysis. Histograms confirm antigen expression in each Raji variant used for co-culture. **D**, Reference UMAP of all viable CD22^+^ tumor events colored by CD19, CD20, or CD38 expression at baseline. **E**, Tumor-cell distribution with AdCAR-T cells alone. **F**, Tumor-cell distribution during individual-AM targeting, colored by the corresponding antigen. **G**, Tumor-cell distribution during combined CD19, CD20, and CD38 targeting. Color scales indicate relative antigen expression from low (blue) to high (red). The flow-cytometric co-culture datasets in panels C–G comprised three technical replicates per condition and time point. Abbreviations: AdCAR, Adapter Chimeric Antigen Receptor; AM, adapter molecule; E:T, effector-to-target ratio; EC₅₀, half-maximal effective concentration; KO, knockout; LLE, linker-label epitope; LLE-tafa, LLE-tafasitamab; LLE-ritu, LLE-rituximab; LLE-dara, LLE-daratumumab; LLE-comb., combination of all three AMs; UMAP, Uniform Manifold Approximation and Projection; WT, wild type.

CD19, CD20, and CD38 represent complementary targets across B-lineage malignancies. CD19 and CD20 are established targets in B-cell leukemias and lymphomas, whereas CD38 extends coverage to plasma-cell malignancies and subsets of aggressive lymphoma (17–21). These antigens therefore constitute a rational target set for broad pan-B-lineage targeting. As CD38 is expressed on activated T cells, CD38-directed targeting may affect the AdCAR-T cell compartment, consistent with reports of fratricide among CD38-specific CAR-T cells (18,22).

The binders used to generate AMs vary considerably in their degree of prior characterization. Most AMs developed to date for the AdCAR-T platform and related adaptor systems rely on *de novo* or research-grade binders. Here, we instead generated CD19- and CD38-directed AMs from in-house-produced tafasitamab and daratumumab biosimilars, respectively, and a CD20-directed AM from commercially available rituximab (19–21). The corresponding therapeutic antibodies are clinically established, providing well-characterized target specificities and clinical safety profiles. Their biotinylation and use within the AdCAR platform nevertheless require AM-specific pharmacological and safety evaluation. Building AMs on clinically validated antibodies is thus a conceptual strategy to accelerate the translation of multi-antigen AdCAR-T targeting.

In this proof-of-concept study, we used a defined antigen-heterogeneous Burkitt lymphoma model comprising wild-type Raji cells and single-antigen knockout variants to investigate individual and combinatorial AM-mediated AdCAR-T targeting. We assessed tumor-cell composition *in vitro*, therapeutic efficacy *in vivo*, and the phenotype and cytotoxic function of AdCAR-T cells following CD38-directed AM exposure. We hypothesized that combinatorial AM targeting would counteract antigen-loss-driven escape without compromising AdCAR-T cell function, supporting antibody-derived AMs as a flexible and translatable pan-B-lineage targeting strategy to address pre-existing antigen heterogeneity.

## Results

### CD19-, CD20-, and CD38-directed AMs mediate antigen-specific, dose-dependent AdCAR-T cell cytotoxicity

To characterize dose-dependent activity and antigen specificity of AdCAR-T cells, we used an in-house-produced, biotinylated tafasitamab biosimilar (LLE-tafasitamab), commercially sourced and biotinylated rituximab (LLE-rituximab), and an in-house-produced, biotinylated daratumumab biosimilar (LLE-daratumumab) as AMs. To test serum stability of the LLE-tag, all three AMs were incubated for 4 or 24 h in native or heat-inactivated human serum. Anti-biotin staining was comparable between conditions for LLE-rituximab and LLE-daratumumab, whereas LLE-tafasitamab showed slightly reduced anti-biotin binding after 24 h in native human serum (Supplementary Figure 1A). Overall, these findings support stability of the AM-biotin configuration under the tested *in vitro* conditions. Next, AdCAR-T cells were co-cultured with CD19^+^CD20^+^CD38^+^ Raji cells (Raji^WT^). All three AMs mediated potent, dose-dependent cytotoxicity, with EC50 values in the low ng/mL range (0.46 ng/mL for LLE-tafasitamab, 0.80 ng/mL for LLE-rituximab, and 0.26 ng/mL for LLE-daratumumab) and more than 90% target-cell lysis at concentrations above 10 ng/mL (Figure 1B).

To confirm antigen specificity, we generated Raji variants, each with a knockout of CD19, CD20, or CD38 (Figure 1C). AdCAR-T cells efficiently lysed Raji^WT^ cells, whereas activity was minimal against the knockout variant lacking the antigen recognized by the respective AM. Each knockout variant remained susceptible to AdCAR-T cell killing through AMs directed against the retained antigens (Supplementary Figure 1B). Thus, cytotoxicity was strictly dependent on the presence of both the AM and its corresponding target antigen, confirming antigen-specific and modular redirection of AdCAR-T cells.

### Combined AM targeting counteracts selection of antigen-deficient tumor subpopulations *in vitro*

To evaluate AdCAR-T cell cytotoxicity in a heterogeneous setting, we generated Raji^KOmix^, an equal mixture of Raji^WT^, Raji^CD19KO^, Raji^CD20KO^, and Raji^CD38KO^ cells. Tumor cell composition during co-culture with AdCAR-T cells was assessed by conventional flow cytometry and UMAP visualization (Figure 1C–G). Without AMs, the relative abundance of the four tumor populations was largely maintained (Figure 1E). Control experiments confirmed that AM concentrations used *in vitro* did not prevent antigen detection by flow cytometry (Supplementary Figure 1C). AdCAR-T cell targeting of individual antigens progressively depleted antigen-positive Raji cells, resulting in enrichment of the corresponding antigen-knockout population among residual tumor cells (Figure 1F; Supplementary Figure 1D). In contrast, simultaneous LLE-tafasitamab, LLE-rituximab, and LLE-daratumumab treatment produced broad depletion across all four subsets, leaving only sparse residual viable tumor events at 72 hours (Figure 1G). These data show that combinatorial targeting prevents selective outgrowth of antigen-deficient populations that emerges under single-antigen targeting.

### Combinatorial targeting of CD19, CD20, and CD38 controls antigen-heterogeneous Burkitt lymphoma *in vivo*

To evaluate AdCAR-T cell activity in an antigen-heterogeneous tumor setting *in vivo*, NSG mice were engrafted with Raji^KOmix^ cells and treated with AdCAR-T cells four days after tumor-cell injection, followed by repeated administration of the indicated AMs (Figure 2A). AdCAR-T cells administered without AMs did not control tumor growth (Figure 2B). Similarly, mice receiving LLE-tafasitamab, LLE-rituximab, or LLE-daratumumab as single AMs progressed to predefined endpoint criteria by day 14. At day 12, the last time point at which all groups remained evaluable, none of the individual-AM groups differed significantly from the AdCAR-only group (adjusted P > 0.9999 for each comparison; Figure 2B, C).

**Figure 2.**
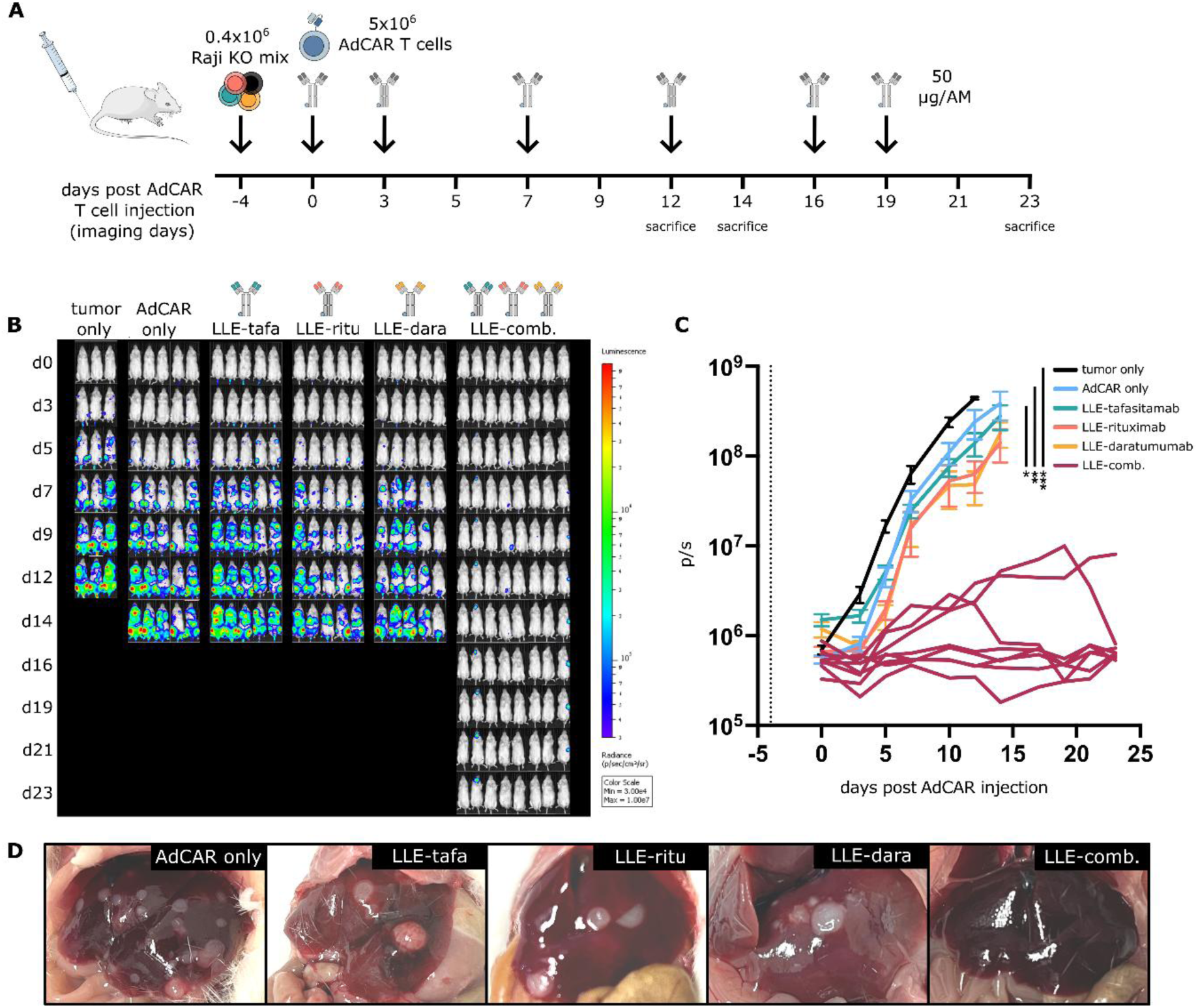
Combined CD19-, CD20-, and CD38-directed AdCAR-T cell targeting controls antigen-heterogeneous Burkitt lymphoma *in vivo*. **A**, NSG mice received 0.4 × 10⁶ Raji^KOmix^ cells intravenously on day −4 and 5 × 10⁶ AdCAR-T cells on day 0, followed by subcutaneous AM administration at the indicated time points (50 µg per AM per dose). Sacrifice time points are indicated in the experimental timeline. **B**, Serial bioluminescence images of mice receiving tumor only (n = 3), AdCAR-T cells without AM (n = 5), LLE-tafasitamab, LLE-rituximab, or LLE-daratumumab (n = 5 each), or all three AMs (LLE-comb.; n = 8). Mice were euthanized after reaching predefined endpoint criteria. **C**, Longitudinal total flux (photons/s). Group means ± SEM are shown for all groups except LLE-comb., for which individual animals are plotted. Day-12 total flux was compared across groups by Kruskal–Wallis test with Dunn’s correction; asterisks indicate significance of each group relative to LLE-comb. (tumor-only, ***; AdCAR-only, **; LLE-tafasitamab, *); comparisons of LLE-comb. to LLE-rituximab and LLE-daratumumab did not reach significance (ns, not shown). **D**, Representative macroscopic liver images from the indicated groups at endpoint. Abbreviations: AdCAR, Adapter Chimeric Antigen Receptor; AM, adapter molecule; LLE-tafa, LLE-tafasitamab; LLE-ritu, LLE-rituximab; LLE-dara, LLE-daratumumab; LLE-comb., combination of all three AMs; ns, P > 0.05; *, P < 0.05; **, P < 0.01; ***, P < 0.001.

In contrast, simultaneous administration of all three AMs resulted in sustained tumor control in 7 of 8 mice throughout the observation period, while one mouse developed a localized head-and-neck signal (Figure 2B, C). At day 12, total flux was significantly lower in the triple-AM group than in the tumor-only (adjusted P = 0.0008), AdCAR-only (adjusted P = 0.0017) and LLE-tafasitamab (adjusted P = 0.0218) groups. Differences compared with the LLE-rituximab (adjusted P = 0.3742) and LLE-daratumumab (adjusted P = 0.8257) groups did not reach statistical significance. Nevertheless, sustained tumor control throughout the observation period was observed only with the triple-AM regimen, whereas all individual-AM groups subsequently progressed to predefined endpoint criteria. Consistent with the bioluminescence data, macroscopic liver examination revealed visible metastases in the AdCAR-only and individual-AM groups, but not in the triple-AM group (Figure 2D). Together, these findings demonstrate sustained tumor control by combinatorial CD19-, CD20-, and CD38-directed targeting in this defined antigen-heterogeneous lymphoma model.

### CD38-directed AM exposure selectively depletes CD38^high^ AdCAR-T cells while preserving a functional CD38^low^ population

Given the expression of CD38 on activated T cells, we next investigated the impact of CD38-directed AM exposure on activated AdCAR-T cells. Following *in vitro* co-culture with Raji^KOmix^ cells, CD38, but not CD19 or CD20, was detected on AdCAR-T cells. Subsequent exposure to LLE-daratumumab resulted in selective loss of the detectable CD38^high^ AdCAR-T cell population, consistent with CD38-dependent fratricide (Figure 3A).

**Figure 3.**
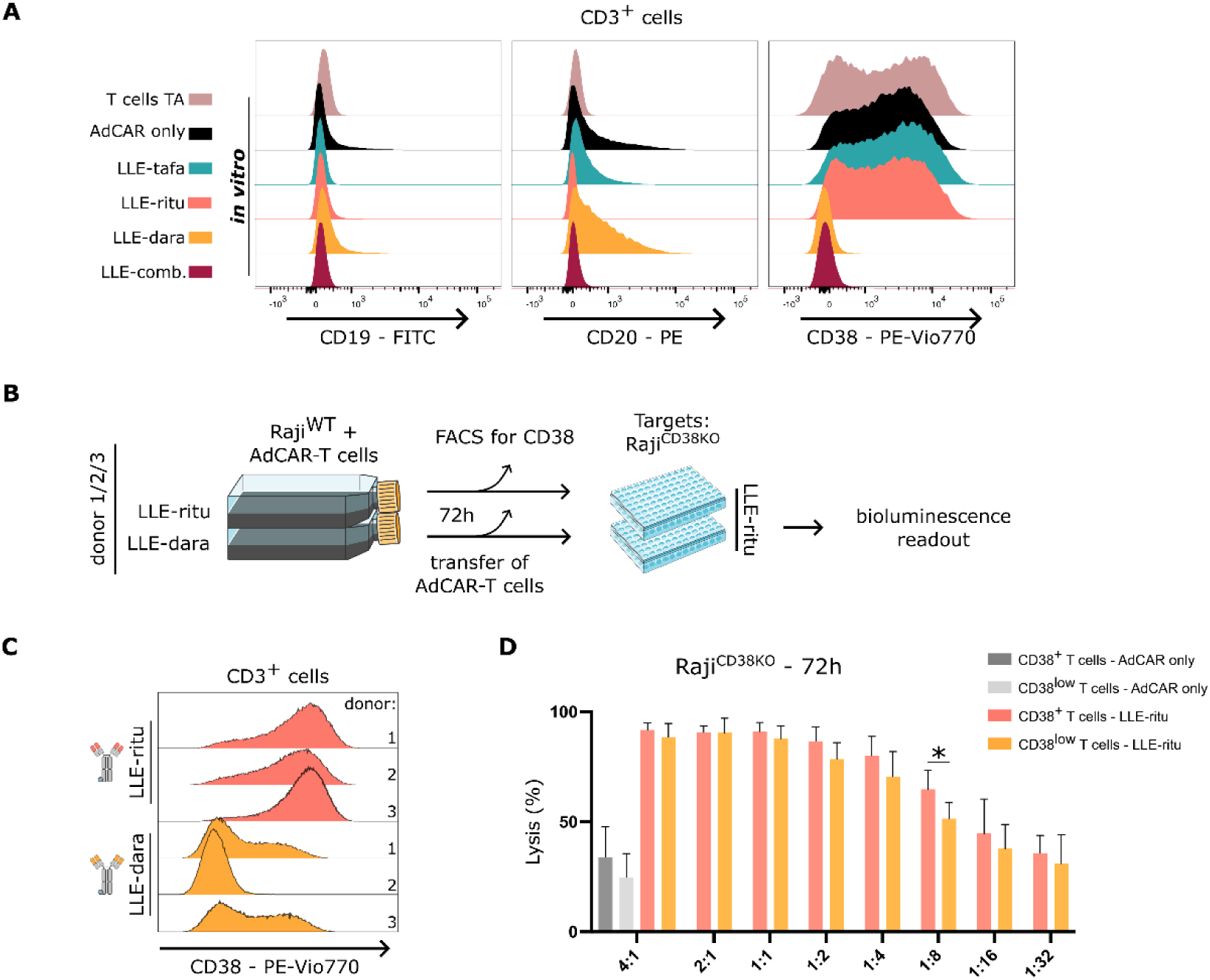
CD38-directed AM exposure selectively depletes CD38^high^ AdCAR-T cells while preserving a functional CD38^low^ population. **A**, CD19, CD20, and CD38 expression on viable CD3^+^ cells. A TransAct-activated T cell sample (T cells TA) is shown as a reference; all other rows show AdCAR-T cells after 72-hour co-culture with Raji^KOmix^ in the absence of AM (AdCAR only) or in the presence of LLE-tafasitamab, LLE-rituximab, LLE-daratumumab, or all three AMs combined (LLE-comb.), each at 10 ng/mL. **B**, Workflow used to generate and functionally assess surviving CD38^low^ AdCAR-T cells. AdCAR-T cells from three donors were co-cultured with Raji^WT^ cells and either LLE-rituximab or LLE-daratumumab (10 ng/ml) for 72 hours (preconditioning). After assessment of CD38 expression, the preconditioned cells were challenged with Raji^CD38KO^ cells using LLE-rituximab as AM. **C**, CD38 expression on AdCAR-T cells from the three donors after preconditioning (see B) with LLE-daratumumab. **D**, Lysis of Raji^CD38KO^ cells after 72 hours at the indicated E:T ratios with 10 ng/mL LLE-rituximab, comparing CD38^+^ AdCAR-T cells (LLE-rituximab-preconditioned) and CD38^low^ AdCAR-T cells (LLE-daratumumab-preconditioned); gray bars show background lysis by AdCAR-T cells without AMs (AdCAR only) for each population. Each donor was tested in three technical replicates, which were averaged within donors for statistical analysis; bars show mean ± SD across the three donor means. Group differences were assessed by one-way ANOVA with Tukey’s multiple-comparisons test; only the comparison between CD38+ and CD38low AdCAR-T cells at the 1:8 E:T ratio reached significance (*, P < 0.05). Abbreviations: AdCAR, Adapter Chimeric Antigen Receptor; E:T, effector-to-target ratio; KO, knockout; LLE-ritu, LLE-rituximab; LLE-dara, LLE-daratumumab; LLE-comb., combination of all three AMs; TA, TransAct; ns, P > 0.05; *, P < 0.05.

To determine whether the surviving AdCAR-T cells retained cytotoxic function, cells were preconditioned for 72 hours by co-culture with Raji^WT^ cells in the presence of either LLE-daratumumab or LLE-rituximab (Figure 3B). Flow cytometric analysis after 72 hours showed marked enrichment of CD3^+^ T cells and no detectable residual CD22^+^ Raji^WT^ population (Supplementary Figure 3). LLE-daratumumab-preconditioned AdCAR-T cells displayed a CD38^low^ phenotype (Figure 3C). The remaining AdCAR-T cells were subsequently challenged with Raji^CD38KO^ cells in the presence of LLE-rituximab, thereby excluding a direct contribution from residual CD38-directed targeting to the cytotoxicity assay.

Both preconditioned AdCAR-T cell populations mediated potent target cell lysis across the tested E:T ratios. CD38^low^ AdCAR-T cells preconditioned with LLE-daratumumab retained cytotoxic activity comparable to that of the LLE-rituximab-preconditioned control at most E:T ratios, with a significant reduction observed at an E:T ratio of 1:8 (adjusted P = 0.021; Figure 3D).

Together, these data demonstrate selective loss of CD38^high^ AdCAR-T cells following CD38-directed AM exposure, consistent with CD38-dependent fratricide, while the surviving CD38^low^ population retained potent cytotoxic activity.

Collectively, these findings demonstrate that combinatorial AdCAR-T cell redirection using AMs derived from clinically validated antibodies can effectively control a defined antigen-heterogeneous lymphoma population *in vitro* and *in vivo*. Notably, CD38-directed AM exposure selectively depleted CD38^high^ AdCAR-T cells, consistent with fratricide, while preserving a CD38^low^ population with sustained cytotoxic capacity.

## Discussion

Antigen downregulation or loss remains a key mechanism of immune escape following CAR-T cell therapy (4–8). Multi-antigen targeting strategies can broaden tumor recognition and reduce antigen-driven escape, but typically rely on target combinations predefined before CAR-T cell infusion and, in some formats, require the manufacture of multiple cellular products (9–11). By separating antigen recognition from CAR signaling, the AdCAR platform offers a modular alternative that enables a single engineered T cell product to be redirected through individual or combined AMs(12–15). Previous work established the feasibility of combinatorial targeting with AdCAR-T cells (14). The present study extends this concept to simultaneous targeting of CD19, CD20, and CD38 in a defined antigen-heterogeneous lymphoma model.

The Raji^KOmix^ system was deliberately reductionist, comprising equal proportions of wild-type cells and three single-antigen knockout variants to generate defined, pre-existing subpopulations and enable direct tracking of antigen-dependent selection over time. AdCAR-T cells together with individual AMs depleted antigen-positive cells while selectively enriching the corresponding antigen-deficient population, whereas triple-AM targeting eliminated all four populations *in vitro*. Consistent with these findings, tumors progressed in all individual-AM groups *in vivo*, while combinatorial targeting achieved sustained tumor control. As dual-AM combinations were not evaluated, these experiments do not define the minimum number of targets required for effective tumor control. Rather, they demonstrate that simultaneous targeting of antigens covering all modeled subpopulations can prevent the predictable outgrowth of antigen-deficient cells under selective pressure.

The use of AMs derived from clinically established antibody frameworks provides a translational advantage, as tafasitamab, rituximab, and daratumumab have well-characterized pharmacological and safety profiles in B-cell and plasma-cell malignancies (19–21). Although in-house production and biotinylation generate distinct investigational reagents that require AM-specific evaluation of dosing, pharmacology, and safety, the underlying antibody frameworks have clinical precedent. Biotin conjugation of the AMs remained stable after 24 hours of incubation in human serum, with anti-biotin antibody binding comparable to that observed in heat-inactivated serum for all AMs except LLE-tafasitamab. LLE-tafasitamab showed slightly reduced anti-biotin antibody binding, suggesting that serum exposure may affect the stability or accessibility of its biotin conjugate. Further optimization of LLE conjugation and AM stability therefore represents an important step in the continued preclinical and translational development of the AdCAR platform.

CD38 targeting highlights a potential liability of AM-mediated redirection, as CD38 expression on activated T cells may render AdCAR-T cells susceptible to fratricide in the presence of CD38-specific AMs. Consistent with this mechanism, LLE-daratumumab exposure selectively depleted the CD38^high^ AdCAR-T cell population. Epitope masking is unlikely to explain this observation, as CD38 remained detectable on Raji^WT^ cells exposed to the same concentration of LLE-daratumumab (10 ng/mL) in the absence of AdCAR-T cells. Together with previous evidence of AM-dependent fratricide following activation-induced CD276 expression in AdCAR-T cells, these findings suggest fratricide as the most likely mechanism underlying the observed population shift (15). However, because T-cell death was not directly quantified, contributions from CD38 internalization or downregulation cannot be excluded. Importantly, the surviving CD38^low^ AdCAR-T cell population retained cytotoxic activity, with significantly reduced activity relative to the control observed only at the lowest E:T ratio (1:8).

CD38 expression has been associated with both activated and dysfunctional T cell states (17,18,22). Whether CD38-directed fratricide preferentially eliminates a functionally distinct AdCAR-T cell subset remains to be determined. Moreover, while the 72-hour rechallenge assay demonstrates retained cytotoxic activity, it did not assess long-term proliferative capacity or persistence. Further studies are therefore needed to determine the functional consequences of CD38-dependent selection within the AdCAR-T cell compartment.

Beyond B-cell malignancies, CAR-T cell-mediated B-cell depletion has recently emerged as a promising therapeutic strategy for severe autoimmune diseases, with CD19-directed CAR-T cells inducing deep B-cell depletion followed by B-cell reconstitution and sustained clinical responses in early clinical studies (23–25). In this setting, the modularity of the AdCAR platform may offer an additional advantage by enabling transient targeting across different stages of the B-cell lineage through the selection and combination of appropriate AMs. Such an approach could potentially achieve broad B-cell depletion while allowing therapeutic pressure to be withdrawn by discontinuing AM administration, warranting further investigation of AdCAR-T cells in B-cell-driven autoimmune diseases.

Combinatorial AdCAR-T cell targeting of CD19, CD20, and CD38 using AMs derived from clinically established antibodies effectively controlled pre-existing antigen heterogeneity and counteracted the outgrowth of antigen-deficient tumor populations. Despite the selective loss of CD38^high^ AdCAR-T cells following CD38-directed AM exposure, the remaining CD38^low^ population retained cytotoxic activity. These findings support further development of the AdCAR-T cell platform as a flexible pan-B-lineage targeting strategy to mitigate antigen-driven tumor escape in B-cell malignancies.

## Methods

### Raji cell culture and reporter-expressing knockout variants

Raji cells (ATCC CCL-86) were maintained in RPMI 1640 (Sigma-Aldrich) supplemented with 2 mM L-glutamine, 100 U/mL penicillin, 100 µg/mL streptomycin, and 10% heat-inactivated fetal bovine serum (Thermo Fisher Scientific) at 37 °C and 5% CO₂. Raji^WT^ and in-house-generated single-antigen knockout variants lacking CD19, CD20, or CD38 were used. Antigen-negative populations were enriched by two consecutive rounds of magnetic cell sorting (MACS), and loss of the respective surface antigen was verified by flow cytometry. All variants expressed a luciferase/mCherry reporter for bioluminescence-based assays and were mixed in equal proportions to generate Raji^KOmix^. Cell cultures were routinely tested and confirmed negative for mycoplasma contamination.

### Production of tafasitamab and daratumumab biosimilars, biotin conjugation of monoclonal antibodies, and serum-stability assessment

Tafasitamab and daratumumab biosimilars were produced in-house in ExpiCHO™ cells (Thermo Fisher Scientific) using the manufacturer’s high-titer transfection protocol. Clarified, filtered culture supernatant was purified by protein A affinity chromatography using GraviTrap columns (Cytiva), followed by desalting and buffer exchange into PBS with PD-10 Sephadex G-25 columns (Cytiva). Commercial rituximab was obtained from the local pharmacy. Antibody concentrations were determined by absorbance at 280 nm using a NanoDrop spectrophotometer (Thermo Fisher Scientific). Each monoclonal antibody was biotinylated with a threefold molar excess of biotin-LC-LC-NHS ester (Thermo Fisher Scientific), following the established laboratory procedure (13,17). Free biotin was removed and buffer was exchanged into PBS using a PD-10 column. Antigen binding and successful biotinylation were verified by flow cytometry on antigen-expressing cell lines using a fluorophore-conjugated anti-biotin detection antibody (Miltenyi Biotec). The resulting biotinylated antibodies are referred to as AMs.

Serum stability of AM-associated biotin was evaluated for all three AMs. Human serum was isolated by centrifugation according to standard procedures from whole blood donated by healthy adult volunteers under Ethics Committee approval 761/2015BO2;an aliquot was heat-inactivated at 56 °C for 30 minutes. Each AM was incubated at 1 µg/ml in untreated or heat-inactivated serum for 4 or 24 hours. Following incubation, AMs were applied to Raji^WT^ cells and antigen-bound AM was detected by flow cytometry using a PE-conjugated anti-biotin antibody (Miltenyi Biotec). Fresh commercial antibodies directed against the corresponding antigens served as immediate-staining reference controls, and secondary-staining-only samples defined background staining. Each condition was evaluated in three technical replicates in one experiment.

### Human T cell isolation, lentivirus production, and AdCAR-T cell generation

The AdCAR construct comprised a signal peptide, an anti-LLE-biotin single-chain variable fragment derived from the murine monoclonal antibody clone mBio3, an extracellular spacer/hinge region, a CD8α transmembrane domain, 4-1BB and CD28 costimulatory domains, and a CD3ζ signaling domain. Lentiviral particles encoding the AdCAR construct were generated by transient transfection of HEK293T cells using polyethyleneimine, concentrated by centrifugation (5,380 × g for 24 hours), resuspended in PBS, and stored at −80 °C.

Peripheral blood mononuclear cells were isolated from whole blood of healthy adult donors recruited at the University Children’s Hospital Tuebingen under Ethics Committee approval 761/2015BO2. Cells were separated by density-gradient centrifugation using Biocoll (Biochrom). T cells were enriched with anti-CD4 and anti-CD8 microbeads (Miltenyi Biotec), activated with TransAct™ (Miltenyi Biotec), and cultured in TexMACS medium (Miltenyi Biotec) supplemented with 10 ng/mL IL-7, 5 ng/mL IL-15 (Miltenyi Biotec), 100 U/mL penicillin, and 100 µg/mL streptomycin. During expansion, lactate was monitored and cultures were diluted with freshly supplemented medium to a lactate concentration of 3 mmol/L; all medium added for expansion contained IL-7 and IL-15 at the stated concentrations.

T cells were transduced with lentiviral particles 36 hours after activation at a multiplicity of infection of 10. Transduction was performed in 200 µL medium by spinoculation at 600 × g and 34 °C for 90 minutes, as used in the closely related AdCAR-T workflow (18). Cells remained in the initial volume for a further 4.5 hours before 1.8 mL of cytokine-supplemented medium was added. AdCAR expression was assessed on days 5–7 by flow cytometry using a PE-labeled AdCAR detection reagent (Miltenyi Biotec); the gating strategy is shown in Supplementary Figure 1E. AdCAR-T cells were expanded for 9 days before *in vivo* experiments and for 12 days before *in vitro* experiments. Unless otherwise stated, in vitro assays used AdCAR-T cells generated from a single donor.

### Bioluminescence-based cytotoxicity assay

Raji cells (2 × 10⁴ per well) were seeded in 96-well plates and co-cultured with AdCAR-T cells at the indicated E:T ratios. Unless otherwise specified, AMs were added at 10 ng/mL for each AM. Co-cultures were maintained in RPMI (Sigma-Aldrich) supplemented with 2 mM L-glutamine, 100 U/mL penicillin, 100 µg/mL streptomycin, 10% heat-inactivated fetal bovine serum (Thermo Fisher Scientific), and 4 µg/mL D-luciferin (PerkinElmer) in a final volume of 200 µL per well at 37 °C and 5% CO₂. Bioluminescence was measured after 24, 48, and 72 hours using a Tecan Spark® microplate reader. Medium-only wells served as no-cell background controls, and untreated tumor-containing wells defined 0% lysis. After subtraction of the medium-only background, tumor-cell viability was calculated by normalizing the luminescence of each treatment well to the mean luminescence of untreated tumor-only wells at the corresponding time point. Specific lysis was calculated as 100 × [1 − (RLU treatment − RLU medium only)/(RLU tumor only − RLU medium only)], as used in the related AdCAR-T workflow (18). Standards containing 100%, 75%, 50%, 25%, and 10% of the initially seeded tumor-cell number confirmed a linear relationship between viable cell number and luminescence and were used to derive relative viable tumor-cell abundance. For the concentration-response experiment, AM concentrations from 0.001 to 100 ng/mL were tested after 72 hours at an E:T ratio of 4:1 with six technical replicates per concentration. Concentration-response data were log-transformed and fitted by nonlinear regression in GraphPad Prism 10; the derived absolute half-maximal concentrations are reported as EC₅₀ values.

### Flow cytometry-based cytotoxicity assay

Raji^WT^, Raji^CD19KO^, Raji^CD20KO^, and Raji^CD38KO^ cells were mixed in equal proportions (0.5 × 10⁵ cells per variant; 2 × 10⁵ total cells) in 5-mL polystyrene round-bottom tubes. AdCAR-T cells were added at an E:T ratio of 4:1 in a final co-culture volume of 1 mL. Unless otherwise stated, each indicated AM was added at 10 ng/mL. Co-cultures were incubated at 37 °C, 5% CO₂, and 95% humidity for the indicated duration. For the longitudinal flow-cytometric co-culture experiment, each experimental condition at each analyzed time point (0, 24, 48, and 72 hours) was established in three technical replicate tubes. Cells were stained in 100 µL CliniMACS buffer (Miltenyi Biotec) for 15 minutes at 4 °C in the dark, washed with 3 mL CliniMACS buffer, and resuspended in 400 µL for acquisition. The panel comprised CD19-FITC (clone LT19, cat. no. 130-113-168; 1:50), CD20-PE (clone REA780, cat. no. 130-111-338; 1:50), CD22-APC-Vio 770 (cat. no. 130-105-060; 1:50), CD38-PE-Vio 770 (clone REA572, cat. no. 130-113-432; 1:50), CD45-APC-Cy7 (clone HI30, cat. no. 304014; 1:50), and CD3-APC or CD3-VioBlue (clone REA613, cat. nos. 130-113-135 and 130-114-519; 1:50 and 1:25, respectively). All antibodies except CD45 (BioLegend) were obtained from Miltenyi Biotec; viability was assessed with 7-AAD (Miltenyi Biotec). Potential interference of surface-bound AM with target-antigen detection was assessed by incubating Raji^WT^ cells for 24 hours with 0, 0.1, 1, 10, 50, 100, or 200 ng/mL LLE-tafasitamab, LLE-rituximab, or LLE-daratumumab before staining for CD19, CD20, or CD38, respectively. The gating strategy is shown in Supplementary Figure 1F. Samples were acquired on a BD FACSCanto™ II using BD FACSDiva™ software (BD Biosciences) and analyzed with FlowJo version 10.8. For visualization of tumor-population composition, samples were downsampled to a common event count before concatenation, and viable CD22^+^ tumor cells from all conditions were embedded together in FlowJo 10.8 using the default UMAP settings, with CD19, CD20, and CD38 as input parameters.

### Animal experiments and *in vivo* procedures

Male 6-to 8-week-old NOD.Cg-Prkdc^scid^ Il2rg^tm1Wjl^/SzJ (NSG) mice were bred and maintained in the animal husbandry facilities of the University Hospital Tuebingen and housed in individually ventilated cages with no more than five animals per cage. General health was monitored daily. All procedures were performed in accordance with the guidelines of the Federation of European Laboratory Animal Science Associations and the approved animal protocol K07/19G. On day −4, mice were injected intravenously with 0.4 × 10⁶ luciferase/mCherry-expressing Raji^KOmix^ cells, comprising a 1:1:1:1 mixture of Raji^WT^, Raji^CD19KO^, Raji^CD20KO^, and Raji^CD38KO^ cells. After randomization, 5 × 10⁶ AdCAR-T cells were administered intravenously on day 0 (gating strategy in Supplementary Figure 2). Beginning on day 0, AMs were administered subcutaneously in 100 µL at alternating 3-and 4-day intervals. Each monotherapy dose contained 50 µg of the indicated AM, whereas the combination dose contained 50 µg of each AM (150 µg total). Each injection additionally contained 10 mg Privigen in the same 100 µL formulation to reduce Fc receptor–mediated AM sequestration. Tumor progression was monitored by bioluminescence imaging at the indicated time points.

### *In vivo* bioluminescence imaging

Mice were anesthetized with isoflurane using a RAS-4 Rodent Anesthesia System (PerkinElmer) and injected subcutaneously with 10 µL/g body weight IVISbrite™ D-luciferin (15 mg/mL in nuclease-free water). Animals were positioned in an IVIS® Lumina Series III imaging chamber (PerkinElmer) with integrated gas anesthesia and a heated stage. After 10 minutes, bioluminescence was recorded at four exposure settings (automatic, 1, 10, and 20 seconds) and analyzed using Living Image® 4.8 software (PerkinElmer). A whole-body region of interest was applied to each animal, and tumor burden was quantified as total flux (photons/s). Mice were euthanized when predefined humane endpoint criteria in the approved animal protocol were reached.

### Cytotoxicity after CD38-directed fratricide

For each of three donors, AdCAR-T cells and Raji^WT^ cells were distributed into parallel T75 flasks at identical cell numbers and co-cultured for 72 hours at an E:T ratio of 4:1 in fully supplemented RPMI containing 10 ng/mL LLE-rituximab or 10 ng/mL LLE-daratumumab. CD38 expression was assessed by flow cytometry to characterize selective loss of the CD38^high^ population after LLE-daratumumab exposure. The designation CD38^low^ was used descriptively for the surviving population with reduced detectable surface CD38; no formal CD38^low^ sorting gate was applied. Flow cytometry demonstrated marked enrichment of CD3^+^ T cells and no detectable residual CD22^+^ Raji^WT^ population after 72 hours (Supplementary Figure 3); therefore, no additional T cell purification was performed. The surviving AdCAR-T cells were counted and evaluated against Raji^CD38KO^ cells at E:T ratios from 4:1 to 1:32 in the bioluminescence-based cytotoxicity assay with 10 ng/mL LLE-rituximab. Each donor-condition pair was tested in three technical replicates, and lysis was assessed after 72 hours.

### Statistical analysis

For the day-12 analysis shown in Figure 2C, total-flux measurements obtained on day 12 were used for all mice. Group differences in day-12 tumor burden were assessed using a Kruskal–Wallis test followed by Dunn’s multiple-comparisons test. In vitro bioluminescence assays were analyzed by one-way ANOVA with Tukey’s multiple-comparisons test; concentration-response curves were fitted by nonlinear regression of log-transformed AM concentrations. For the CD38^low^ assay, technical replicates were averaged within each donor, and the three donors constituted the statistical units. Matched LLE-rituximab- and LLE-daratumumab-preconditioned donor means were compared at each E:T ratio using paired, two-tailed t tests with Bonferroni adjustment across E:T ratios. Analyses were performed in GraphPad Prism 10. Adjusted P values < 0.05 were considered statistically significant; ns, P > 0.05; *, P < 0.05; **, P < 0.01; ***, P < 0.001.

## Abbreviations

7-AAD: 7-aminoactinomycin D
AdCAR: Adapter Chimeric Antigen Receptor
AdCAR-T: Adapter Chimeric Antigen Receptor T cell
AML: acute myeloid leukemia
AM: adapter molecule
ANOVA: analysis of variance
APC: allophycocyanin
ATCC: American Type Culture Collection
BLI: bioluminescence imaging
CAR: chimeric antigen receptor
CD: cluster of differentiation
Cy7: cyanine 7
DKFZ: German Cancer Research Center
DKTK: German Cancer Consortium
E:T: effector-to-target ratio
EC₅₀: half-maximal effective concentration
Fc: fragment crystallizable
FITC: fluorescein isothiocyanate
iFIT: Image-Guided and Functionally Instructed Tumor Therapies
IL: interleukin
IVIS: in vivo imaging system
KiTZ: Hopp Children’s Cancer Center Heidelberg
KO: knockout
LC: long chain
LLE: linker-label epitope
LLE-comb.: combination of all three adapter molecules
LLE-dara: LLE-daratumumab
LLE-ritu: LLE-rituximab
LLE-tafa: LLE-tafasitamab
MACS: magnetic cell sorting
NHS: N-hydroxysuccinimide
NSG: NOD scid gamma
PBS: phosphate-buffered saline
PE: phycoerythrin
RLU: relative luminescence units
RPMI: Roswell Park Memorial Institute
SD: standard deviation
SEM: standard error of the mean
TA: TransAct
UMAP: Uniform Manifold Approximation and Projection
WT: wild type

## Acknowledgements

The authors gratefully acknowledge Kathrin Wolsing and Aysegül Canak for technical assistance.

## Author contributions

SK, DA, and CMS conceived and designed the study. CMS supervised the project. SK, ASM, SS, BK, and DA performed the experiments and analyzed and interpreted the data. SK drafted the manuscript with contributions from ASM, SS, BK, and CMS. All authors reviewed and approved the final manuscript.

## Data availability

The datasets generated and/or analyzed during the current study are available from the corresponding author upon reasonable request.

## References

1. Maude SL, Laetsch TW, Buechner J, Rives S, Boyer M, Bittencourt H, et al. Tisagenlecleucel in Children and Young Adults with B-Cell Lymphoblastic Leukemia. N Engl J Med. 2018 Feb;378(5):439–48. doi:10.1056/NEJMoa1709866

2. Neelapu SS, Locke FL, Bartlett NL, Lekakis LJ, Miklos DB, Jacobson CA, et al. Axicabtagene Ciloleucel CAR T-Cell Therapy in Refractory Large B-Cell Lymphoma. N Engl J Med. 2017 Dec 28;377(26):2531–44. doi:10.1056/NEJMoa1707447

3. Schuster SJ, Bishop MR, Tam CS, Waller EK, Borchmann P, McGuirk JP, et al. Tisagenlecleucel in Adult Relapsed or Refractory Diffuse Large B-Cell Lymphoma. N Engl J Med. 2019 Jan 3;380(1):45–56. doi:10.1056/NEJMoa1804980

4. Sotillo E, Barrett DM, Black KL, Bagashev A, Oldridge D, Wu G, et al. Convergence of Acquired Mutations and Alternative Splicing of *CD19* Enables Resistance to CART-19 Immunotherapy. Cancer Discov. 2015 Dec 1;5(12):1282–95. doi:10.1158/2159-8290.CD-15-1020

5. Orlando EJ, Han X, Tribouley C, Wood PA, Leary RJ, Riester M, et al. Genetic mechanisms of target antigen loss in CAR19 therapy of acute lymphoblastic leukemia. Nat Med. 2018 Oct;24(10):1504–6. doi:10.1038/s41591-018-0146-z

6. Plaks V, Rossi JM, Chou J, Wang L, Poddar S, Han G, et al. CD19 target evasion as a mechanism of relapse in large B-cell lymphoma treated with axicabtagene ciloleucel. Blood. 2021 Sep 23;138(12):1081–5. doi:10.1182/blood.2021010930

7. Duell J, Leipold AM, Appenzeller S, Fuhr V, Rauert-Wunderlich H, Da Via M, et al. Sequential antigen loss and branching evolution in lymphoma after CD19- and CD20-targeted T-cell–redirecting therapy. Blood. 2024 Feb 22;143(8):685–96. doi:10.1182/blood.2023021672

8. Majzner RG, Mackall CL. Tumor Antigen Escape from CAR T-cell Therapy. Cancer Discov. 2018 Oct 1;8(10):1219–26. doi:10.1158/2159-8290.CD-18-0442

9. Fousek K, Watanabe J, Joseph SK, George A, An X, Byrd TT, et al. CAR T-cells that target acute B-lineage leukemia irrespective of CD19 expression. Leukemia. 2021 Jan;35(1):75–89. doi:10.1038/s41375-020-0792-2

10. Wang T, Tang Y, Cai J, Wan X, Hu S, Lu X, et al. Coadministration of CD19- and CD22-Directed Chimeric Antigen Receptor T-Cell Therapy in Childhood B-Cell Acute Lymphoblastic Leukemia: A Single-Arm, Multicenter, Phase II Trial. J Clin Oncol. 2023 Mar 20;41(9):1670–83. doi:10.1200/JCO.22.01214

11. Spiegel JY, Patel S, Muffly L, Hossain NM, Oak J, Baird JH, et al. CAR T cells with dual targeting of CD19 and CD22 in adult patients with recurrent or refractory B cell malignancies: a phase 1 trial. Nat Med. 2021 Aug;27(8):1419–31. doi:10.1038/s41591-021-01436-0

12. Seitz CM, Mittelstaet J, Atar D, Hau J, Reiter S, Illi C, et al. Novel adapter CAR-T cell technology for precisely controllable multiplex cancer targeting. OncoImmunology. 2021 Jan;10(1):2003532. doi:10.1080/2162402X.2021.2003532

13. Atar D, Mast AS, Scheuermann S, Ruoff L, Seitz CM, Schlegel P. Adapter CAR T Cell Therapy for the Treatment of B-Lineage Lymphomas. Biomedicines. 2022 Sep 28;10(10):2420. doi:10.3390/biomedicines10102420

14. Atar D, Ruoff L, Mast AS, Krost S, Moustafa-Oglou M, Scheuermann S, et al. Rational combinatorial targeting by adapter CAR-T-cells (AdCAR-T) prevents antigen escape in acute myeloid leukemia. Leukemia. 2024 Oct;38(10):2183–95. doi:10.1038/s41375-024-02351-2

15. Kristmann B, Werchau N, Suresh L, Pezzuto EL, Scheuermann S, Krost S, et al. Targeting CD276 with Adapter-CAR T-cells provides a novel therapeutic strategy in small cell lung cancer and prevents CD276-dependent fratricide. J Hematol OncolJ Hematol Oncol. 2025 Jul 28;18(1):76. doi:10.1186/s13045-025-01729-8

16. Volta L, Myburgh R, Pellegrino C, Koch C, Maurer M, Manfredi F, et al. Efficient combinatorial adaptor-mediated targeting of acute myeloid leukemia with CAR T-cells. Leukemia. 2024 Dec;38(12):2598–613. doi:10.1038/s41375-024-02409-1

17. Calabretta E, Carlo-Stella C. The Many Facets of CD38 in Lymphoma: From Tumor– Microenvironment Cell Interactions to Acquired Resistance to Immunotherapy. Cells. 2020 Mar 26;9(4):802. doi:10.3390/cells9040802

18. Li W, Liang L, Liao Q, Li Y, Zhou Y. CD38: An important regulator of T cell function. Biomed Pharmacother. 2022 Sep;153:113395. doi:10.1016/j.biopha.2022.113395

19. Lonial S, Weiss BM, Usmani SZ, Singhal S, Chari A, Bahlis NJ, et al. Daratumumab monotherapy in patients with treatment-refractory multiple myeloma (SIRIUS): an open-label, randomised, phase 2 trial. The Lancet. 2016 Apr;387(10027):1551–60. doi:10.1016/S0140-6736(15)01120-4

20. Duell J, Abrisqueta P, Andre M, Gaidano G, Gonzales-Barca E, Jurczak W, et al. Tafasitamab for patients with relapsed or refractory diffuse large B-cell lymphoma: final 5-year efficacy and safety findings in the phase II L-MIND study. Haematologica. 2023 Aug 31;109(2):553–66. doi:10.3324/haematol.2023.283480

21. Maloney D, Liles T, Czerwinski D, Waldichuk C, Rosenberg J, Grillo-Lopez A, et al. Phase I clinical trial using escalating single-dose infusion of chimeric anti-CD20 monoclonal antibody (IDEC-C2B8) in patients with recurrent B-cell lymphoma. Blood. 1994 Oct 15;84(8):2457–66. doi:10.1182/blood.V84.8.2457.2457

22. Piedra-Quintero ZL, Wilson Z, Nava P, Guerau-de-Arellano M. CD38: An Immunomodulatory Molecule in Inflammation and Autoimmunity. Front Immunol. 2020 Nov 30;11:597959. doi:10.3389/fimmu.2020.597959

23. Mackensen A, Müller F, Mougiakakos D, Böltz S, Wilhelm A, Aigner M, et al. Anti-CD19 CAR T cell therapy for refractory systemic lupus erythematosus. Nat Med. 2022 Oct;28(10):2124–32. doi:10.1038/s41591-022-02017-5

24. Müller F, Taubmann J, Bucci L, Wilhelm A, Bergmann C, Völkl S, et al. CD19 CAR T-Cell Therapy in Autoimmune Disease — A Case Series with Follow-up. N Engl J Med. 2024 Feb 22;390(8):687–700. doi:10.1056/NEJMoa2308917

25. Schett G, Müller F, Taubmann J, Mackensen A, Wang W, Furie RA, et al. Advancements and challenges in CAR T cell therapy in autoimmune diseases. Nat Rev Rheumatol. 2024 Sep;20(9):531–44. doi:10.1038/s41584-024-01139-z

