## Supplementary Material for "Pan-B-Lineage Targeting with Adapter CAR-T Cells Controls Antigen-Heterogeneous Lymphoma"

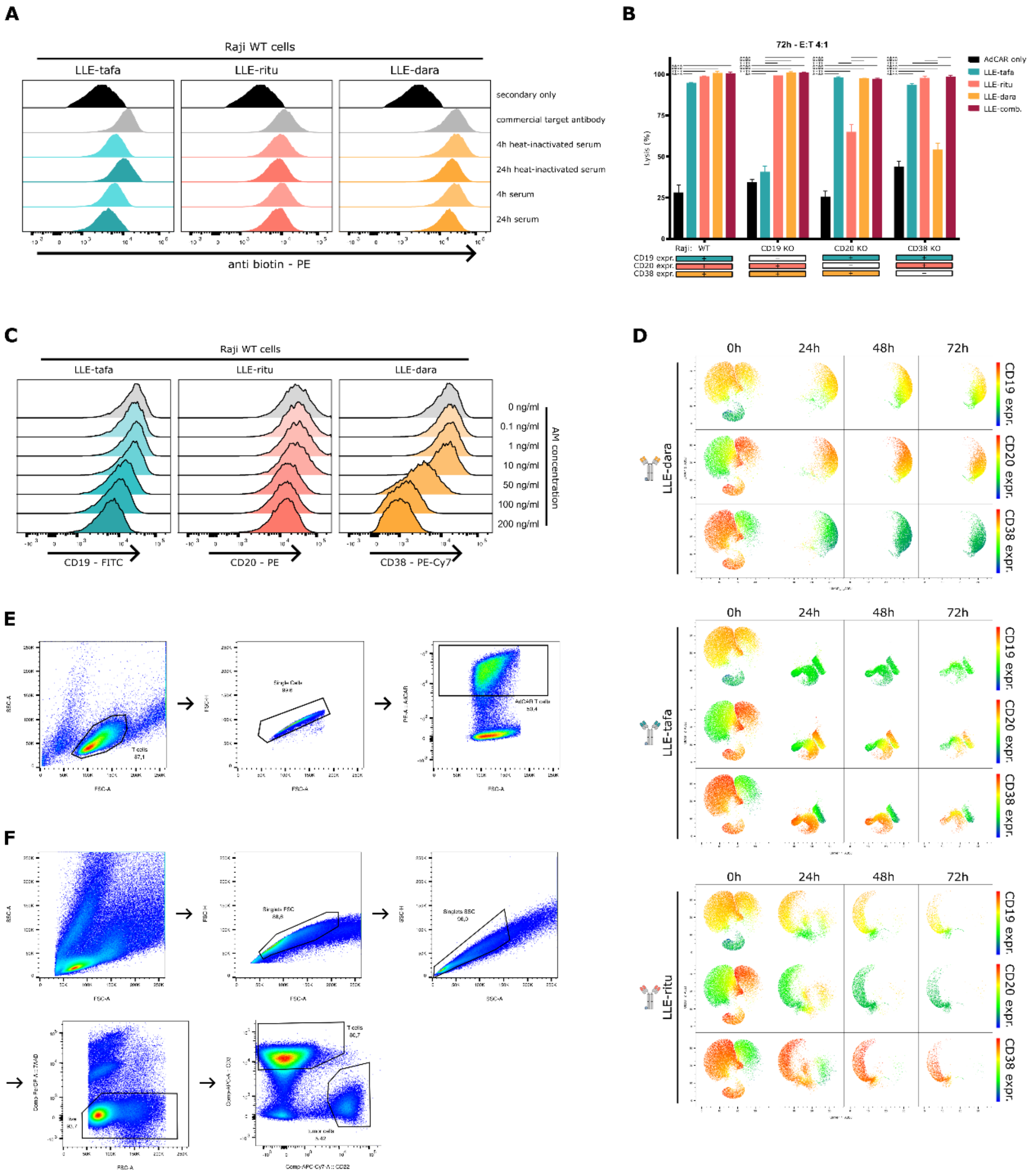


**Supplementary Figure 1. Serum stability of the LLE-tag and flow-cytometric controls for *in vitro* AdCAR-T cell assays.**

**A**, Serum stability of the LLE-tag after incubation of LLE-tafasitamab, LLE-rituximab, and LLE-daratumumab for 4 or 24 hours in native or heat-inactivated human serum. AMs were subsequently incubated with Raji$^{WT}$ cells to allow antigen binding, followed by detection of AM-associated biotin using a PE-conjugated anti-biotin antibody. Corresponding commercial antigen-specific antibodies and secondary-only samples served as independent staining controls. Histograms are representative of one experiment with three technical replicates per condition. **B**, Lysis of Raji$^{WT}$, Raji$^{CD19KO}$, Raji$^{CD20KO}$, and Raji$^{CD38KO}$ cells after 72-hour co-culture with AdCAR-T cells at an E:T ratio of 4:1 and the indicated AMs. Bars show mean ± SD. C, Antigen detection on Raji$^{WT}$ cells after 24-hour incubation with the indicated concentrations of LLE-tafasitamab, LLE-rituximab, or LLE-daratumumab. At 10 ng/mL, the concentration used in the *in vitro* assays, the corresponding antigens remained detectable by flow cytometry. **D**, UMAP visualization of viable tumor cells during individual targeting with LLE-tafasitamab, LLE-rituximab, or LLE-daratumumab at 0, 24, 48, and 72 hours; each condition and time point was analyzed in three technical replicates. Events are colored by expression of the indicated antigen from low (blue) to high (red). **E**, Flow-cytometric gating strategy used to identify AdCAR-expressing T cells with the AdCAR detection reagent. **F**, Gating strategy used to identify viable T cells and tumor cells in the flow cytometry-based cytotoxicity assay. Abbreviations: AdCAR, Adapter Chimeric Antigen Receptor; AM, adapter molecule; E:T, effector-to-target ratio; KO, knockout; LLE, linker-label epitope; LLE-tafa, LLE-tafasitamab; LLE-ritu, LLE-rituximab; LLE-dara, LLE-daratumumab; LLE-comb., combination of all three AMs; UMAP, Uniform Manifold Approximation and Projection; WT, wild type; ns, P > 0.05; *, P < 0.05; **, P < 0.01; ***, P < 0.001.

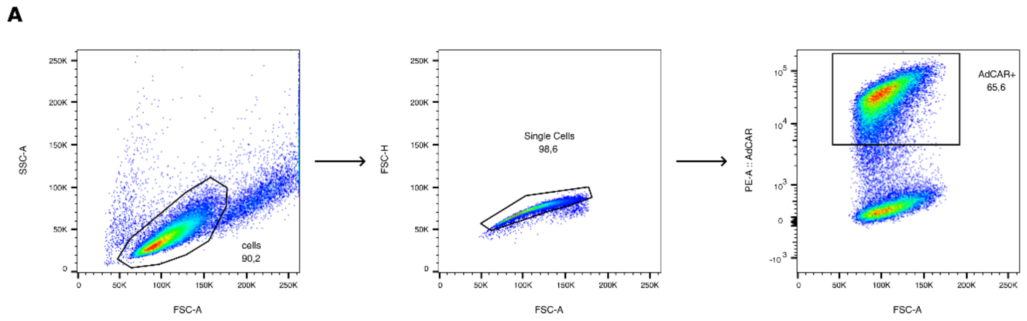


**Supplementary Figure 2. Pre-infusion flow-cytometric gating of AdCAR-T cells.**
Representative gating strategy used to quantify AdCAR expression on AdCAR-T cells before *in vivo* administration.

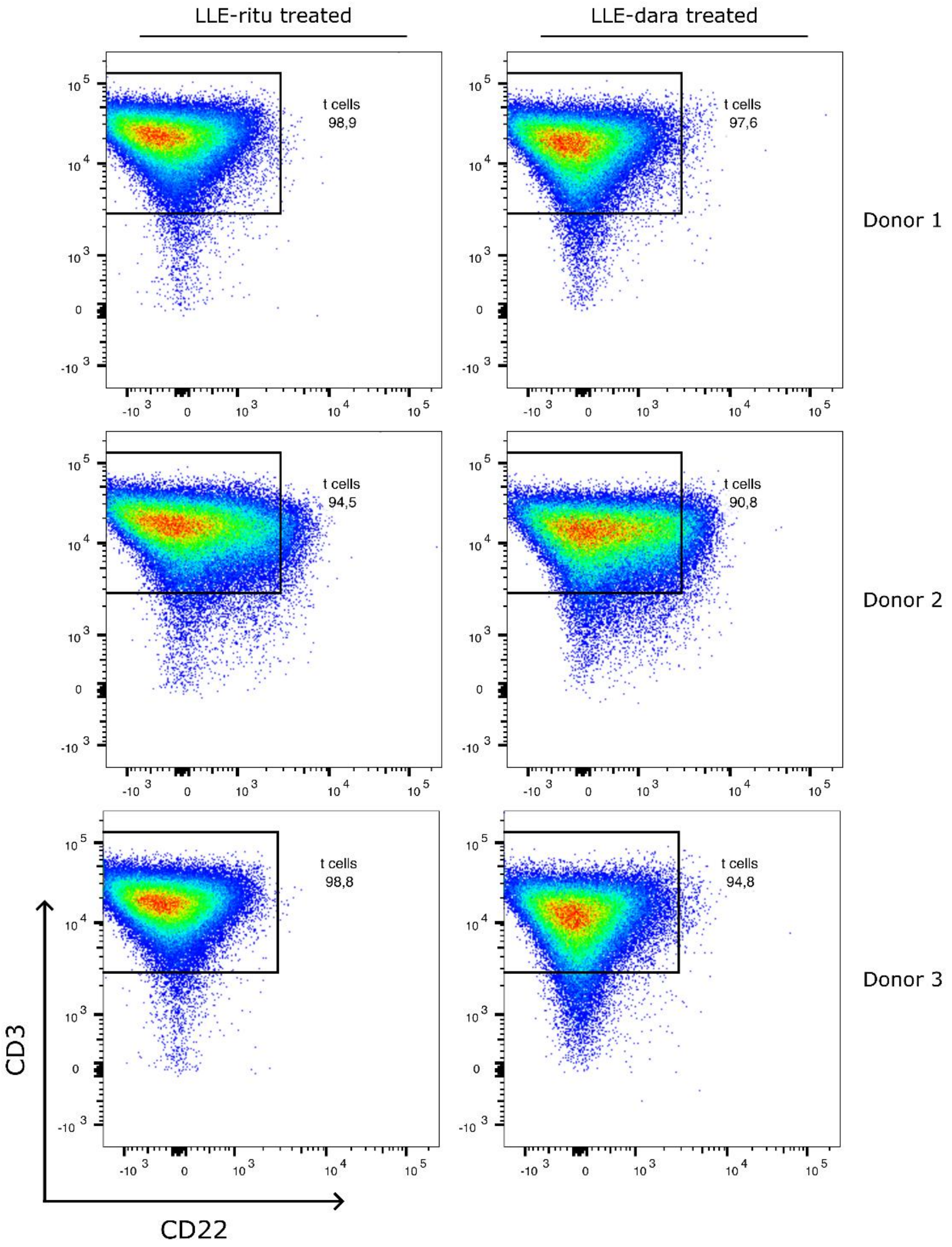


**Supplementary Figure 3. Flow-cytometric confirmation of T cell enrichment after AM-mediated preconditioning.**

CD3 and CD22 expression after 72-hour preconditioning with Raji$^{WT}$ cells and LLE-rituximab (left) or LLE-daratumumab (right) for three donors. Percentages indicate CD3$^{+}$ T cells among living single cells.
